# Cytokine-induced Chromatin Accessibility in Whole Blood Neutrophils Links to Sepsis Transcriptional States

**DOI:** 10.64898/2026.02.03.703332

**Authors:** Justin Cayford, Brandi Atteberry, Akanksha Singh-Taylor, Andrew Retter, Benjamin P. Berman, Theresa K Kelly

## Abstract

**Background:** Neutrophils play an important role in the immune system and sense environmental perturbations including pathogens. Upon pathogen detection, neutrophils extrude their chromatin, forming neutrophil extracellular traps (NETs) trapping and removing pathogens. Previous studies have shown that controlled chromatin decondensation occurs during NET formation, reflecting NET inducing pathways, and the cellular environment. While NET inducing stimuli like phorbol 12-myristate 13-acetate (PMA) is commonly used to study NET formation, it bypassing regulatory mechanisms, limiting insights.

**Methods:** We used the Assay for Transposase-Accessible Chromatin with sequencing (ATAC-Seq) to profile chromatin accessibility in neutrophils stimulated in whole blood with PMA and physiologically relevant inflammatory factors (NFs), including TNF-α, GM-CSF, fMLP, C5a, and IL-1β, alone and in combination. Chromatin responses were compared across conditions and integrated with publicly available transcriptomic sepsis cohorts.

**Results:** We found that NF stimulation induced stimulus specific chromatin accessibility programs distinct from PMA. Individual NFs increased specific transcription factor (TF) motif enrichments in a stimulus dependent manner, with GM-CSF increasing STATs, TNF-α increasing NF-κB, C5a/fMLP increasing AP-1, and Combined with a cooperative response including CEBP. Integration with sepsis transcriptomic datasets revealed that promoter accessibility changes within NF stimulations correspond to transcriptional states associated with sepsis disease severity, highlighting the upstream regulatory programs linked to clinical outcomes.

**Conclusions:** These findings demonstrate that NF stimulation in whole blood reveals chromatin accessibility programs in neutrophils that correlate with disease severity in sepsis. This approach provides a framework for linking cytokine driven neutrophil regulation to heterogenous inflammatory states in sepsis and other NET-associated diseases.

## 1. Introduction

Neutrophils are central to the innate immune system and can form neutrophil extracellular traps (NETs), web-like structures of decondensed chromatin and antimicrobial proteins designed to immobilize and kill microbes (1-3). While NET formation contributes to host defense, excessive or dysregulated NET release is linked to immunothrombosis, vascular injury, organ dysfunction, and increased mortality (4-8). These maladaptive effects are particularly evident in NET associated inflammatory diseases such as sepsis, which was associated with ∼11 million deaths worldwide in 2017 (9).

Despite extensive evidence implicating NETs in disease pathology, the mechanisms underlying individual sensitivity to excessive NET formation and immune dysregulation remain poorly understood (10-12). Clinical outcomes in sepsis vary widely even among patients with similar infectious triggers, suggesting that immune regulatory mechanisms contribute to disease heterogeneity (10, 11, 13-15). Large transcriptomic studies have demonstrated that sepsis comprises multiple immune endotypes rather than a uniform inflammatory state (13, 14, 16, 17). However, the regulatory mechanisms that shape neutrophil transcriptional programs and responses to NET inducing stimuli remain poorly defined.

Chromatin accessibility is a crucial regulatory layer that precedes and constrains transcription by controlling transcription factor (TF) binding (18, 19). Profiling chromatin accessibility provides a powerful framework for linking inflammatory signaling to functional immune states. However, neutrophils present unique challenges for epigenomic profiling due to their short lifespan, rapid activation, and sensitivity to isolation protocols (20).

To address these limitations, we previously established a fixed ATAC-Seq strategy that preserves chromatin accessibility following stimulation and enables robust profiling of neutrophils directly from whole blood without prior isolation (21, 22). Using this approach, we previously characterized chromatin accessibility changes in neutrophils in isolation and in whole blood stimulated with phorbol 12-myristate 13-acetate (PMA), a known potent inducer of NET formation that acts mainly through protein kinase C (PKC) (18, 21-25). These studies demonstrated that neutrophil chromatin is highly structured, responds in a coordinated manner to stimulation, and exhibits more robust accessibility changes in whole blood consistent with complex immune system signaling (22).

While PMA has been widely used to study NETs, it bypasses many physiological regulatory pathways by directly activating PKC (24, 26). However, *in vivo* neutrophils are exposed to complex mixtures of naturally occurring cytokines and chemokines (natural factors; NFs) that engage distinct signaling pathways (10, 11). In sepsis, circulating levels of tumor necrosis factor-α (TNF-α), granulocyte-macrophage-colony-stimulating factor (GM-CSF), complement component C5a, interleukin-1β (IL-1β), and formylated peptides (fMLP) are often elevated (11, 27-31) and have been implicated in neutrophil priming, activation, and NET formation (32, 33).

Here, we sought to define how physiologically relevant NF stimulation remodels neutrophil chromatin accessibility in whole blood, and whether these accessibility changes correspond to clinically relevant transcriptional states. Building on our previous high-throughput NET formation study identifying cytokine combinations capable of inducing NET release (34), we stimulated healthy whole blood with individual NFs and in combination. We show that NF stimulation engages a broader inflammatory landscape than PMA and that promoter accessibility changes correlate with transcriptional endotypes observed in sepsis cohorts (15-17, 35). Together, these data establish a framework for potentially linking NF driven chromatin regulation in neutrophils to endotypes in sepsis or other NET-associated inflammatory diseases (11, 15-17, 35).

## 2. Materials and Methods

### 2.1 Ethics approval

Whole blood was obtained from healthy donors in K2 EDTA tubes (BD #366643; PrecisionMed, San Diego). Research was approved under WCG IRB Protocol number 20181025, and all human participants gave written informed consent. Donors were self-reported healthy, aged 18-50, with BMI < 30, and not taking NSAIDs for at least 24 hours prior to donation (Supplementary Table 1).

### 2.2 Whole blood treatment

Whole blood stimulation with PMA was performed as previously described (22), and expanded to include naturally occurring factors (NFs). Briefly, 2 mL of pooled whole blood was aliquoted into 5 mL tubes and stimulated with PMA (250 nM), TNF-α (10 µg/mL), GM-CSF (10 µg/mL), fMLP (10 µg/mL), C5a (10 µg/mL), IL-1β (85 ng/mL), a combination of all NFs (Combo), or vehicle controls at 37°C for 120 minutes with inversion every 30 minutes (Supplementary Table 2).

Whole blood was fixed with a 10x formaldehyde solution (Sigma #252549), 1M NaCl, 0.1 mM EDTA (Fisher Scientific #AM9010), and 0.5 mM HEPES (ThermoFisher #15630080) for 10 minutes at room temperature and quenched with 2.5M glycine. Fixed neutrophils were isolated using the MACSxpress Whole Blood Neutrophil Isolation Kit (Miltenyi Biotec) and ATAC-Seq was performed on 125,000 neutrophils per sample.

### 2.3 ATAC-Seq library preparation and processing

ATAC-Seq libraries were generated using recombinant Tn5 transposase (Active Motif #81284) as previously described (19, 21, 22, 36) and sequenced on an Illumina NovaSeq platform (Active Motif). Data were processed using the nf-core/atacseq Nextflow pipeline (v2.1.2) (https://nf-co.re/atacseq/2.1.2) (37) with alignment to the hg38 reference genome. Peaks were called using an FDR<0.01, and ENCODE quality standards were applied (38) (Supplementary Figure 1, Supplementary Table 3).

To generate overlapping peak sets, peaks detected in at least one technical replicate were retained and combined across donors. Peaks present in at least two-thirds of donors were retained for downstream analysis. Variance partitioning was performed using a linear mixed-effects model to estimate the relative contributions of stimulation, donor, and residual variance (39).

### 2.4 Differential accessibility analysis

Differentially accessible regions (DARs) were identified using DESeq2 (28) as previously described (22) . Briefly, peaks with a baseMean>10, adjusted p-value<0.01, and |log_2_(fold change)|>1 were retained for downstream analysis. Principal Component Analysis (PCA) was performed using prcomp, heatmaps were generated using pheatmap (https://cran.r-project.org/web/packages/pheatmap/index.html). UpSetR plots (https://cran.r-project.org/web/packages/UpSetR/index.html) were generated using the .boolean.annotatePeaks.txt output files.

### 2.5 Enrichment analysis

Motif enrichment analysis was performed using HOMER (40) (findMotifsGenome.pl, hg38) with default parameters (22). Background regions were generated by random sampling from the full set of ATAC peaks. Odds ratios (ORs) were calculated as:

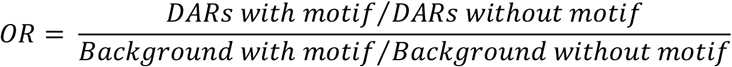

Odds ratios were used as effect-size measures rather than for statistical inference. Motifs were grouped into transcription factor (TF) families for comparative analysis. Promoter associated DARs were identified using annotatePeaks.pl (40) and resulting promoter peaks were retained.

Genomic Regions Enrichment of Annotations Tool (GREAT) (41, 42) was used for functional interpretation. BED files were uploaded with default parameters and whole genome background was used. GREAT’s binomial test with FDR correction was used and terms with FDR<0.05 were retained.

### 2.6 Clustering analysis of DARs

DARs from all vehicle-vs-stimulus comparisons, excluding PMA, were combined for unsupervised clustering. A composite significance score was calculated per region and comparison as:

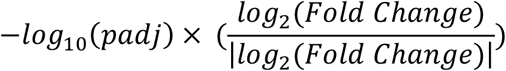

Hierarchical clustering was performed using Euclidean distance and Ward’s linkage (ward.D2) with the number of clusters (k=10) was selected using elbow and silhouette analyses (Supplementary Figure 3B). Cluster level accessibility profiles were calculated as the mean composite score per cluster and stimulus and z-score normalized for visualization.

### 2.7 VANISH and SUBSPACE transcriptomic cohort integration and ROC analysis

The VANISH cohort (35) was used to define neutrophil-associated transcriptional modules (CTS1-CTS3) (15) and calculate per-gene mean expression values across donors. Genes with overlapping promoter DARs (pDARs) were assigned to CTS modules based on relative expression patterns.

Gene expression data from the SUBSPACE cohort (15) were extracted for genes overlapping pDARs and donors were stratified by annotated clinical severity. Receiver operating characteristic (ROC) analysis was performed using the pROC R package (43) and confidence intervals for area under the ROC curve (AUC) estimates were calculated using bootstrap resampling.

## 3. Results

### 3.1 Stimulation of whole blood with PMA and natural factors induce a neutrophil response

Healthy donor whole blood was treated with PMA, individual NFs (TNF-α, GM-CSF, fMLP, C5a, IL-1β; Supplementary Table 2), or NFs in combination (Combo), followed by fixation, neutrophil isolation, and ATAC-Seq (Figure 1A). Peaks were called across donors to generate an overlapping peak set for each condition (Figure 1B, 1C, Supplementary Figure 2A).

**Figure 1.**
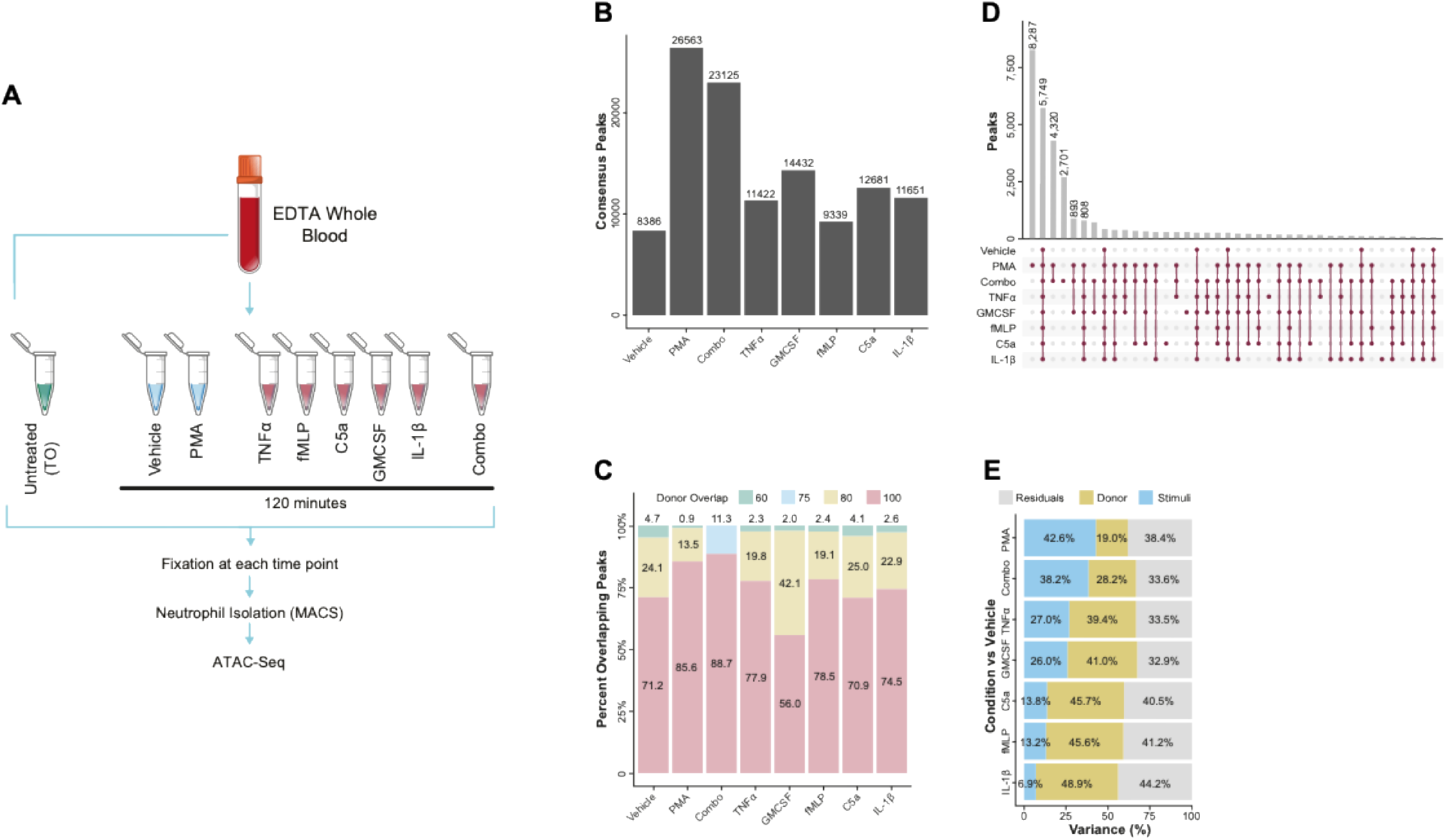
Stimulation of whole blood with PMA and natural cytokines induce neutrophil response: **(A)** Experimental schematic. Whole blood was collected (n=5) and vehicle, 250 nM phorbol 12-myristate 13-acetate (PMA), 10 µg/mL TNF-α, fMLP, C5a, GM-CSF, 85 ng/mL IL-1β, and Combo containing the same concentrations (TNF-α, fMLP, C5a, GM-CSF, and IL-1β; n=4) and incubated for 120 minutes, formaldehyde fixed, neutrophils were isolated, and then ATAC-Seq was completed. **(B)** Total number of consensus peaks across all donors and stimulation conditions, the peak must have been retained in one replicate and then in at least 3 donors (66% of donors), there were n=4 donors for Combo and n=5 donors for all other conditions. **(C)** The percentage of consensus peaks that were shared across each stimulation condition and donor overlap. **(D)** UpSetR plot showing the shared peaks across the stimulation consensus peaks. **(E)** Variance partitioning analysis across the consensus peak set of each individual stimulation (PMA, Combo, TNF-α, GM-CSF, C5a, fMLP, and IL-1β). Blue indicates the stimulus fraction contribution, yellow indicates the fraction contributed by the donor, and grey indicates all residual fraction contributions. The percent is indicated for each category and stimulation.

PMA induced the largest number of peaks (26,563), followed by the Combo (23,125 peaks), whereas individual NFs induced fewer peaks (9,339-14,432) (Figure 1B). The vehicle controls exhibited 8,386 peaks, indicating increased chromatin accessibility across all stimulation conditions (Figure 1B).

Peak overlap was highly reproducible across donors, with all stimuli except GM-CSF showing >70% overlap and both PMA and Combo exceeding 85%. Stronger stimuli appeared to reduce donor-to-donor variability, suggesting that they can partially overcome donor-to-donor variability (Figure 1C).

Analysis of peak overlap revealed 5,789 peaks were shared across all conditions, suggesting a core neutrophil accessibility signature (Figure 1D). PMA induced 8,287 unique peaks, whereas Combo had 2,701 unique peaks, suggesting a distinct regulatory signature not recapitulated by PMA. Despite these differences, PMA and Combo shared 4,320 peaks, indicating a robust shared activation of chromatin. Interestingly, we observed that fMLP was the only treatment condition without a unique accessibility pattern (Figure 1D).

Variance partitioning demonstrated that PMA and Combo accounted for the largest fraction of accessibility variance (43% and 38%, respectively) and were the only stimuli for which stimulation variance overcame donor variance (PMA: 19%, Combo: 28%; Figure 1E). Individual NFs showed greater donor-to-donor variance, suggesting less robust stimulation. Together, these results indicate that weaker stimulation leads to more donor-specific differences in chromatin accessibility and may also reflect specific pathway activation.

### 3.2 PMA and Combo treatments induce a similar, yet distinct neutrophil response

Based on their robust stimulation effect, we next focused on PMA and Combo. PCA revealed clear separation between vehicle, PMA, and Combo, with minimal donor-dependent clustering, indicating stimulus driven remodeling (Figure 2A).

**Figure 2.**
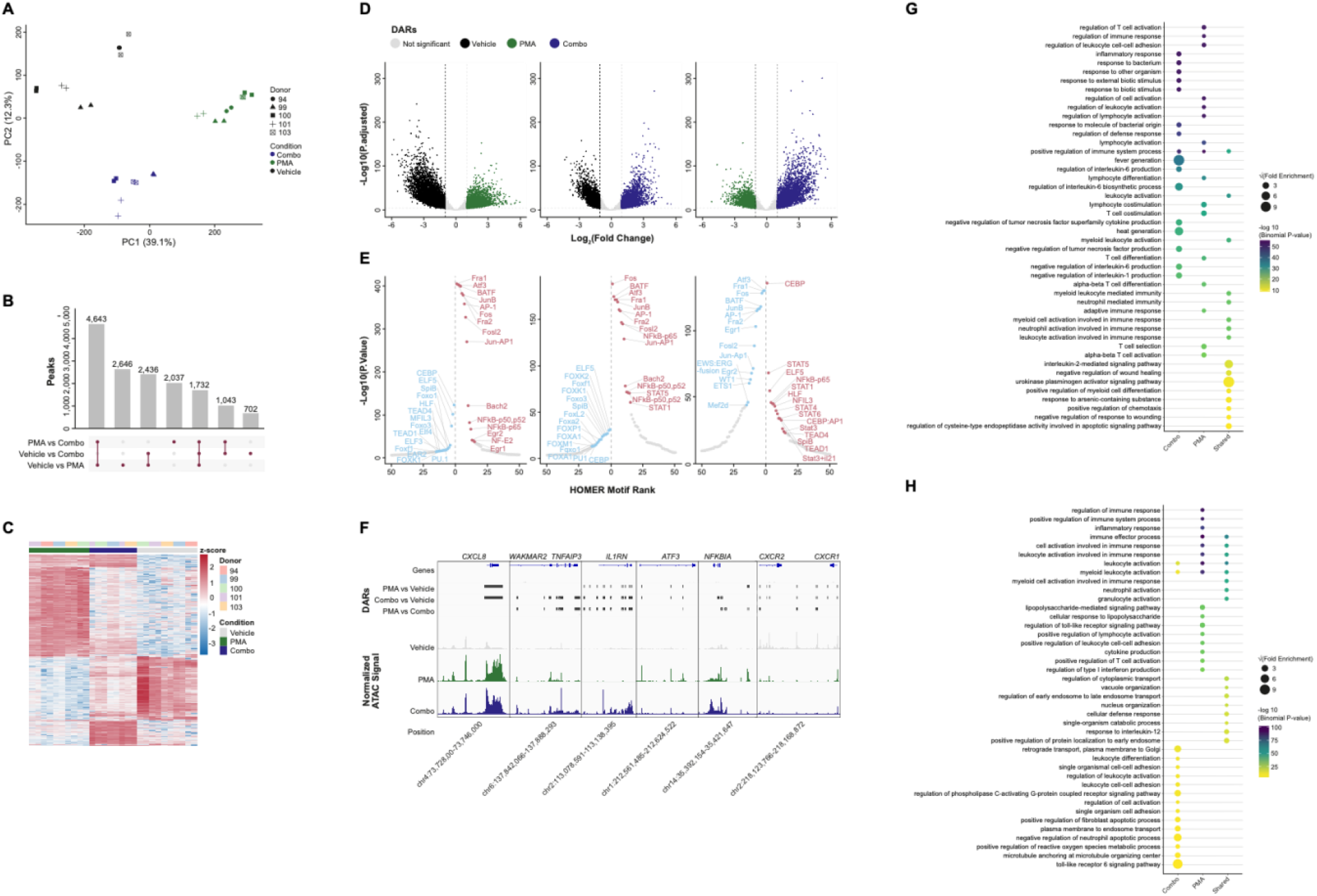
PMA and Combo treatments induce a similar, yet distinct neutrophil response: **(A)** Principal Component Analysis across all donors (donors are indicated by: circle: 94, triangle: 99, square: 100, plus: 101, square with x: 103) and selected stimulation conditions (black: vehicle, green: PMA, blue: Combo). PC1 (x-axis) represents 39.1% of the total variance and PC2 (y-axis) represents 12.3% of variance. Donor numbers are indicated by the number adjacent to the datapoint. **(B)** UpSetR plot showing the overlap of differentially accessible regions (DARs) comparing the three conditions (Vehicle vs PMA, Vehicle vs Combo, and PMA vs Combo). **(C)** Hierarchically clustering of Z-score normalized peak counts across the top 5,000 DARs (sorted based on p.adjusted value) from the three comparisons (grey: Vehicle, green: PMA, blue: Combo) across all donors (pink: 94, blue: 99, green: 100, purple: 101, orange: 103).. **(D)** Volcano plots of DARs in each comparison (Left: vehicle vs PMA, middle: vehicle vs Combo, right: PMA vs Combo) with log2(fold change) (x-axis) and -log10(p.adjusted) (y-axis). The DARs are colored by significance and dashed lines indicate threshold (-log10(p.adjusted) > 4, log2(fold change) > 1.25 or log2(fold change) < -1.25) in each comparison (grey: not significant, black: vehicle, green: PMA, blue: Combo). **(E)** HOMER motifs within the significant DARs in each comparison from (D) (Left: vehicle vs PMA, middle: vehicle vs Combo, right: PMA vs Combo). Results are plotted by -log10(p.value) (y-axis) and HOMER rank (x-axis). Increased motifs are indicated in red (right side of individual panel) and decreased motifs are indicated in blue (left side of individual panel). **(F)** Merged donor tracks (grey: vehicle, green: PMA, blue: Combo) visualized using the IGV genome browser at various loci with normalized ATAC signal. Significant DARs indicated by bars (PMA (dark grey) vs Vehicle (light grey), Combo (dark grey) vs Vehicle (light grey), and PMA (light grey) vs Combo (dark grey)). **(G)** Dot plot showing GREAT pathway analysis of increasing accessible DARs from comparing PMA and Combo to vehicle. These were broken up to unique in Combo (left) or PMA (middle) and shared (right). Dot size indicates the square root of the absolute value log2(fold enrichment) and color indicates the -log10(binomial p value) generated by GREAT. **(H)** Similar to (G) but the pathway analysis was completed using the decreasing accessible DARs from the same comparisons.

Next, we identified differentially accessible regions (DARs) and found a large set of DARs (4,643) shared between PMA-versus-vehicle and PMA-versus-Combo, suggesting PMA accessibility changes that were not recapitulated by Combo (Figure 2B). Combo stimulation produced a smaller unique set of 702 DARs specific to Combo-versus-vehicle, and an additional 1,403 DARs that were in both Combo-versus-vehicle and Combo-versus-PMA (Figure 2B). PMA and Combo shared a common set of DARs (2,436) relative to vehicle, suggesting a similar yet distinct response from the two stimulations (Figure 2B).

Visualization of the top 5,000 of DARs (32.8%) following z-score normalization showed attenuated responses can be seen with distinct subsets of regions between PMA and Combo stimulation (Figure 2C), indicating the varied responses. Volcano plots demonstrated that PMA induced nearly double the number of DARs as Combo (11,457 versus 5,913; Figure 2D; Supplementary Table 4), with PMA showing a large fraction of DARs exhibiting reduced accessibility (n=6,493 decreasing versus n=4,964 increasing) (Figure 2E). In contrast, Combo produced a more balanced pattern of increasing (n=2,815 DARs) and decreasing accessibility (n=3,098 DARs) (Figure 2D). Direct comparison of PMA to Combo highlighted stronger PMA-associated accessibility gains relative to Combo (PMA increasing, n=3,814 DARs; Combo increasing, n=5,641 DARs) (Figure 2D).

Motif enrichment analysis (HOMER) revealed strong enrichment of AP-1 family motifs (JUN/FOS/FRA) in regions with increased accessibility following PMA stimulation, consistent with previous reports (22) (Supplementary Figure 2B; Figure 2ECombo stimulation also enriched AP-1 motifs but showed additional enrichment of STAT (STAT1/3/4/5/6) and NF-κB motifs, consistent with cytokine-mediated inflammatory signaling (Figure 2E). Notably, CEBP motifs were preferentially enriched in Combo specific regions, whereas AP-1 motifs dominated PMA specific regions (Figure 2E), highlighting divergent regulatory mechanisms. Motifs associated with decreased accessibility differed slightly between conditions, with PMA specific motifs having a depletion of ETS, TEAD and FOX family of motifs, consistent with the repression of homeostatic regulatory elements, while Combo showed depletion in the FOX family (Figure 2E).

Representative loci of the dynamic chromatin responses of PMA and Combo were shown in Figure 2F. Both PMA and Combo increased accessibility at the *CXCL8* locus and decreased accessibility at *CXCR1/2*. In contrast, *TNFAIP3* showed greater accessibility following Combo stimulation, and *NFKBIA* showed PMA specific enhancer activation (Figure 2F). Together, these examples illustrate that while PMA and Combo share many DARs and TF motifs, their regulatory pathways can be quite different.

Pathway analysis using GREAT (41, 42) revealed that increased accessibility regions were enriched for immune activation pathways in both conditions. PMA favored broad leukocyte activation pathways, while Combo had enriched cytokine-mediated processes, such as IL-6 and regulation of TNF signaling (Figure 2G). Decreased accessibility regions were also distinct, with PMA repressing immune effector pathways and Combo affecting vesicular trafficking and apoptotic regulation (Figure 2H).

Together, these data suggest PMA and Combo induce overlapping activation signatures while also showing distinct regulation between activation and repression in whole blood stimulated neutrophils.

### 3.3 Individual natural factors induce factor specific chromatin changes

To understand the contribution of individual NFs, we performed UMAP projection across all conditions. Combo clustered most closely with GM-CSF and showed limited donor variability, while TNF-α was clustered near Combo and GM-CSF, but displayed partial donor dependency (Figure 3A). GM-CSF had a donor (100) that clustered far from the group, suggesting a unique donor impact. Other stimuli (C5a, fMLP, IL-1β, and vehicle) clustered more distantly and indicated greater donor variability.

**Figure 3.**
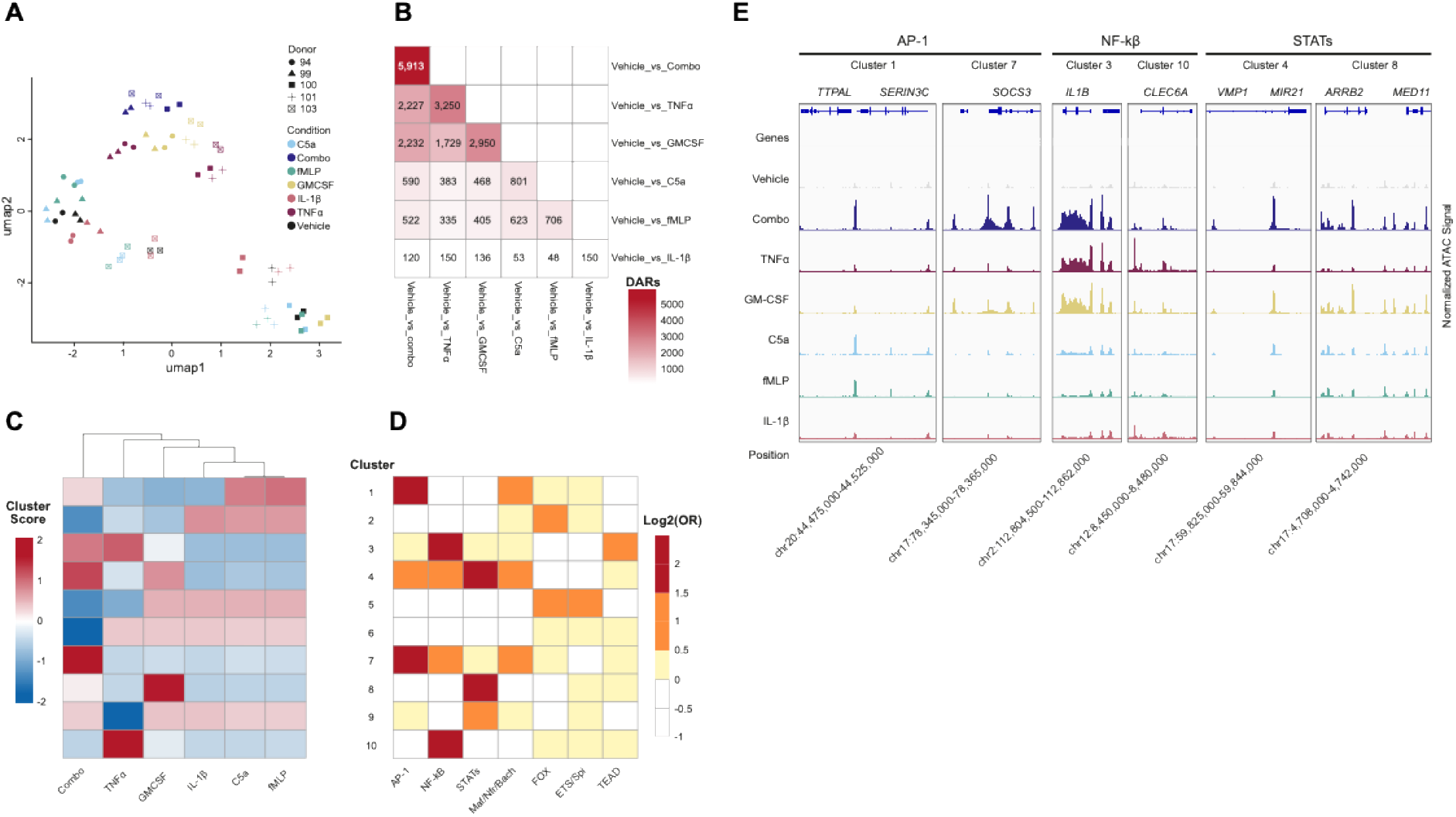
Individual natural factors induce factor specific chromatin changes: **(A)** UMAP projection across natural factor stimulations (grey: vehicle, dark blue: Combo, dark red: TNF-α, yellow: GM-CSF, light blue: C5a, green: fMLP, light red: IL-1B). UMAP1 (x-axis) and UMAP2 (y-axis) are shown. Donors are indicated by circle: 94, triangle: 99, square: 100, plus: 101, square with x: 103. **(B)** Total numbers of DARs overlapping across each stimulation compared to vehicle. Diagonal values indicate the total number of DARs identified in that comparison. **(C)** Unsupervised hierarchical clustering heatmap (k=10) on DAR composite activation score and then Cluster Signal was used to visualize (mean signal across entire cluster) across all stimulations (x-axis) and assigned cluster (y-axis). Columns were hierarchical clustered. **(D)** Heatmap of HOMER Motif analysis across each cluster (y-axis). Motifs were collapsed into the indicated motif family (x-axis). Color indicates increasing fold enrichment from log2(Odds Ratio) (white: -1-0, yellow: 0-0.5, orange: 0.5-1, red: 1.5-3). **(E)** Merged donor tracks (grey: vehicle, dark blue: Combo, dark red: TNF-α, yellow: GM-CSF, light blue: C5a, green: fMLP, light red: IL-1B) visualized using the IGV genome browser at various loci with normalized ATAC signal.

Differential accessibility analysis (DESeq2 (28); FDR<0.01, baseMean>10) identified overlapping DARs across stimulation comparisons and showed the number varied widely by stimulation (Figure 3B; Supplementary Table 5). Combo had the largest number (n=5,913), followed by TNF-α and GM-CSF (3,250 and 2,950, respectively), whereas C5a and fMLP induced fewer than 1,000 DARs and IL-1β yielded a minimal response (n=150) (Figure 3B). Overlap analysis demonstrated both shared and distinct accessibility changes, with Combo and GM-CSF having the largest number of overlaps (2,232/5,913; 38%) followed by TNF-α (2,227/5,913; 37%). C5a and fMLP shared the largest overlap percentage (623/801; 77.8%) despite limited overlap across other stimulation (Figure 3B and Supplementary Figure 3A).

Unsupervised hierarchical clustering (Ward.D2) of DARs across NF conditions identified distinct regulatory modules (Figure 3C; Supplementary Figure 3B). The analysis yielded cluster modules ranging from 215-1,949 DARs (mean≈758 ± 490) (Supplementary Figure 3C). We visualized cluster level mean accessibility patterns, revealing cluster patterns across stimuli (Figure 3C). Motif enrichment analysis revealed clear TF structure across clusters after grouping broad TF families (AP-1, NF-κB, STAT, Maf/Nrf/Bach, FOX, ETS/Spi, and TEAD) for visualization. AP-1 motifs (JUN/FOS/FRA/BATF) dominated clusters associated with Combo and C5a/fMLP, NF-κB motifs were enriched in TNF-α associated clusters, and STAT motifs were enriched in GM-CSF driven clusters (Figure 3D). Maf/Nrf/Bach and weaker FOX/ETS/Spi motifs were weakly observed across clusters and stimulations, which could reflect that the chromatin remodeling was not a primary driver (Figure 3D).

Representative loci illustrated specific regulation within the clusters. AP-1 associated clusters showed selective promoter accessibility at the *TTPAL/SERIN3C* (Cluster 1) and SOCS3 (Cluster 7) loci (Figure 3E). NF-κB-enriched clusters showed increased accessibility at *IL-1β* (Cluster 3) and *CLEC6A* (Cluster 10). STAT-linked clusters showed GM-CSF and Combo driven accessibility at *MIR21* (Cluster 4) and *ARRB2* (Cluster 8) loci (Figure 3E).

Taken together, these data demonstrate that individual NFs activate distinct yet overlapping chromatin regulatory patterns in neutrophils. Combo stimulation integrated multiple TF networks suggesting a broad, physiologic inflammatory response across multiple pathways.

### 3.4 Chromatin accessibility signatures associated with transcriptional programs linked to progression of severe disease

To assess whether NF induced NET chromatin programs reflected transcriptional profiles observed in sepsis, we integrated our results with publicly available transcriptomic datasets. We first focused on the VANISH cohort (44), which had defined three septic transcriptomic states (CTS1-CTS3) associated with clinical and drug treatment outcomes (15).

To increase interpretability and better align with expression datasets, we restricted our analysis to promoter-associated DARs (pDARs) across all NF stimulations and compared these regions to VANISH expression data (Figure 4A). Within each CTS module, genes were classified by concordance between promoter accessibility and transcription (up/up or down/down). Scatterplots of the pDAR accessibility versus VANISH expression highlighted concordant genes within each CTS module (Figure 4A). We found that ∼50% of genes were concordant in each module and the up/up genes were a larger fraction than the down/down (Supplementary Table 6).

**Figure 4.**
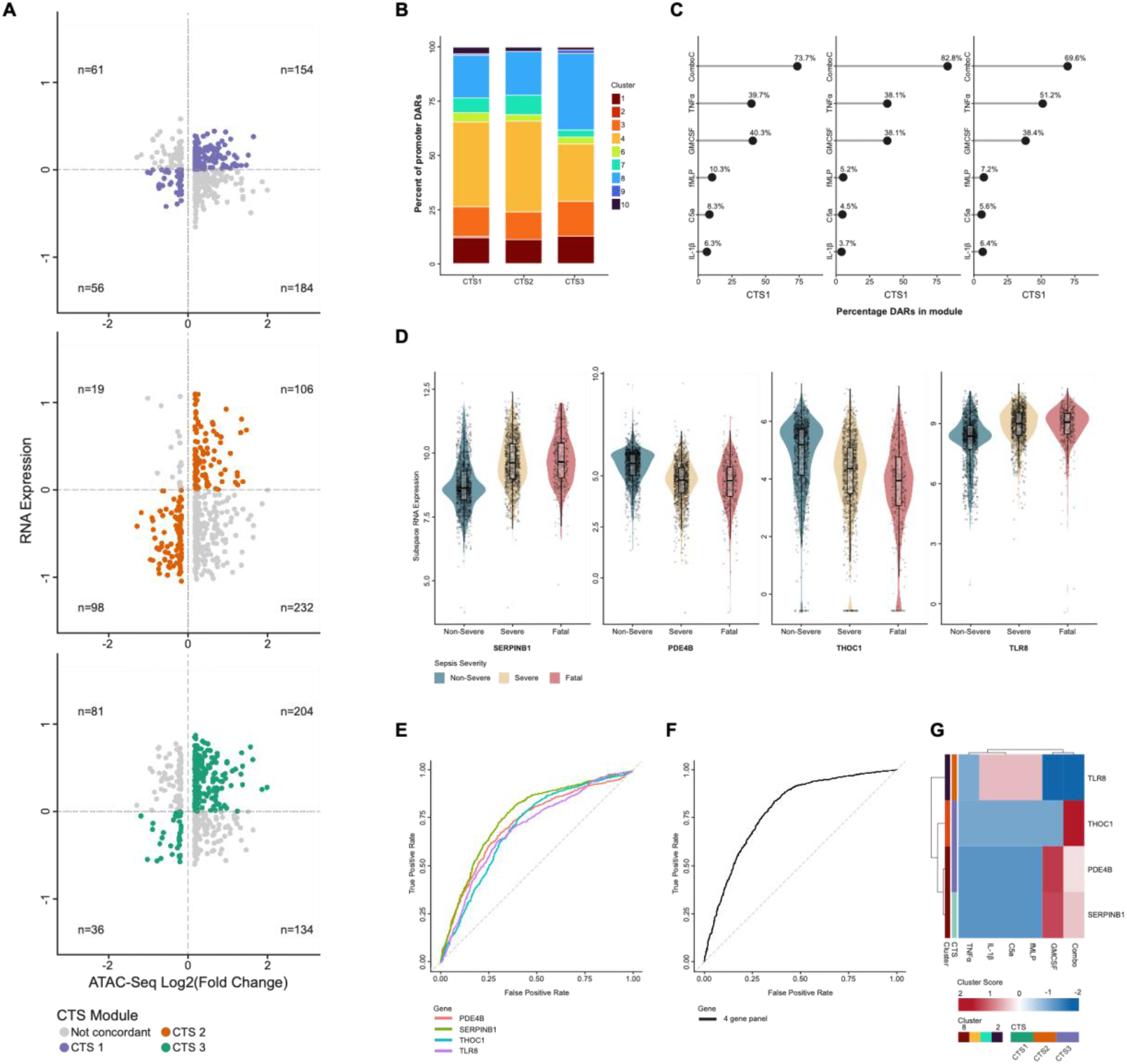
Chromatin accessibility signatures associated with transcriptional programs linked to progression of severe disease: **(A)** Scatterplots comparing promoter-associated chromatin accessibility changes to mean gene expression values (across donors) from the VANISH cohort. Genes are grouped by CTS module and colored by concordant directionality between accessibility and transcription. Top – CTS1; green points indicate concordance – n=204 for up/up and n=36 for down/down. Middle – CTS2; orange points indicate concordance – n=106 for up/up and n=96 for down/down. Bottom – CTS3; purple points indicate concordance – n=154 for up/up and n=56 for down/down. **(B)** Distribution of ATAC cluster assignments among concordant promoter-DARs for each CTS module. Bars represent the proportion of regions assigned to each chromatin cluster, highlighting shared and distinct regulatory composition. For cluster assignment colors: 1 – dark blue, 2 - blue, 3 - light blue, 4 – teal, 5 – green, 6 – light green, 7 – light orange, 8 – orange, 9 – red, 10 – dark red. **(C)** Lollipop plots summarizing the relative contribution of individual stimulation conditions to CTS associated promoter accessibility changes. For each CTS module, the percentage of regions showing differential accessibility is shown. **(D)** Expression of ATAC-linked genes in the SUBSPACE cohort across sepsis severity grades (Green – non-severe, yellow – severe, and red – fatal). Violin and boxplots show the distribution of normalized gene expression for the four genes (*SERPINB1, PDE4B, THOC1*, and *TLR8*) retained for downstream analysis. Individual points represent individual donors (non-severe: n=1,293, severe: n=1,203, and fatal: n=387). **(E)** Receiver operating characteristic (ROC) curves evaluating the ability of genes to distinguish severe/fatal from non-severe disease in the SUBSPACE cohort. Four genes all contained an area under the curve (AUC) > 0.70 (Green - *SERPINB1*; AUC = 0.76, Red - *PDE4B*; AUC = 0.72, Blue - *THOC1*; AUC = 0.7, and Purple - *TLR8*; AUC = 0.7). **(F)** Similar to (E) but ROC curve for all four genes (*SERPINB1, PDE4B, THOC1*, and *TLR8*) combined (AUC=0.79). **(G)** Heatmap of promoter-associated chromatin accessibility for the four genes across all stimulation conditions. Rows represent the promoter-associated peak per gene, and columns represent the stimulation compared to vehicle. Values are z-score normalized across the comparisons. Row annotations indicate ATAC cluster assignment and CTS module (CTS1 – green, CTS2 – orange, CTS3 – purple).

We next asked whether concordant CTS genes mapped to specific ATAC clusters. Using the clusters defined in Figure 3D, we examined cluster representation across CTS groups and found that CTS1 and CTS2 had similar compositions, dominated by clusters 7 and 3 (enriched for AP-1 and NF-κB) (Figure 4B; Supplementary Figure 4A). CTS3 showed a minor shift towards cluster 3 (NF-κB) and reduced representation of the AP-1 dominated clusters (Figure 4B). Despite these minor differences, all CTS modules exhibited a similar balance of AP-1, NF-κB, and STAT motif signatures.

To determine which NF stimuli contributed most to CTS linked pDARs, we attributed each concordant pDAR to the stimulation condition where it was observed. Across CTS modules, accessibility changes were primarily driven by Combo, with the secondary contributions from GM-CSF and TNF-α (Figure 4C). In contrast, C5a, fMLP, and IL-1β accounted for a substantially smaller fraction of CTS-linked regions (Figure 4C; Supplementary Figure 4B). These results suggest that CTS-linked accessibility regions are more closely associated with Combo, TNF-α, and GM-CSF.

To assess whether these genes also associated with a larger, more clinically stratified cohort, we next examined expression in the SUBSPACE cohort (15), which included non-severe, severe, and fatal sepsis. Intersection of VANISH pDARs with SUBSPACE yielded 10 shared genes (Supplementary Figure 4C). Given the limited overlap, we treated this analysis as exploratory and focused on identifying consistent severity linked expression trends. Visualizing expression across the severity grades (excluding healthy controls to focus on progression) indicated several genes changed between the non-severe and severe/fatal groups (Figure 4D; Supplemental Figure 4C). A subset of four genes (*TLR8, THOC1, PDE4B*, and *SERPINB1*) showed the most consistent association with severity (Figure 4D).

To quantify separation of non-severe versus severe/fatal disease, we performed ROC analysis. The four genes achieved AUC values >0.70, indicating modest but reproducible separation of disease progression (Figure 4E). When combining all four genes, the AUC was 0.79, suggesting cooperativity of these genes in disease progression (Figure 4F). Notably, TLR8, THOC1, and PDE4B were mapped to the CTS3 module, while SERPINB1 (highest individual AUC) mapped to CTS1. Because this study was not designed as a diagnostic analysis, these results are interpreted as trends highlighting candidate genes rather than biomarkers.

Finally, to link these candidates back to NF stimulation, we examined chromatin accessibility profiles for the four genes across our treatment conditions. GM-CSF and Combo stimulation showed consistent accessibility changes across these genes, with more variability for TNF-α and minimal effects from C5a, fMLP, and IL-1β (Figure 4G). This pattern was consistent with cluster identity and suggests that accessibility of the genes was dominated by GM-CSF and to a lesser extent TNF-α (Figure 4G, Supplementary Figure 4D).

Collectively, these analyses connect NF induced chromatin remodeling in whole blood neutrophils to transcriptional programs observed in clinical sepsis cohorts. While the observed associations are modest and the clinical integration is exploratory, the consistency across independent datasets supports the relevance of NF promoter accessibility programs to inflammatory states linked to disease progression.

## 4. Discussion

In this study, we demonstrated that physiologically relevant stimulation of whole blood elicited distinct and organized chromatin accessibility responses in neutrophils that integrate multiple inflammatory signaling pathways and correspond to transcriptional states observed in sepsis (11, 15-17, 35). While PMA induced the most extensive chromatin remodeling, NF stimulation produced a more balanced regulatory landscape including CEBP, AP-1, NF-κB, and STAT signaling (24, 25). This distinction is critical, since PMA induces predominantly PKC dependent activation, whereas NF stimulation more closely reflects physiologic inflammatory signaling, including accessibility gains at inflammatory loci and compaction at regulatory elements involved in vesicular trafficking and apoptosis, known processes of NET formation. The selective enrichment of CEBP associated motifs in Combo suggests neutrophil chromatin responses reflect engagement of lineage defining architecture rather than purely acute activation pathways (45, 46).

Clustering of DARs revealed modular regulatory programs associated with specific stimuli in neutrophils. GM-CSF activated STAT clusters, TNF-α activated NF-κB clusters, and C5a/fMLP activated AP-1 clusters (47, 48). Notably, Combo stimulation did not reflect a linear additive effect of the individual NFs, but instead produced a cooperative state dominated by GM-CSF and TNF-α, with additional AP-1 engagement. This suggests nonlinear integration of inflammatory signals in neutrophil chromatin regulation.

The use of whole blood is a key feature of this study. Our prior work demonstrated that neutrophil isolation alters baseline chromatin accessibility with only 57% overlap in accessibility peaks in unstimulated neutrophils when fixed before and after isolation (22). Furthermore, stimulation in whole blood produced more robust chromatin changes and allowed for the potential influence of secondary mediators/intercellular signaling to be captured (22). Extending this approach to NF stimulation reveals regulatory features that are absent or attenuated in isolated systems and are particularly relevant for inflammatory diseases such as sepsis, where neutrophils actively respond to the dynamic environment in disease progression (8, 10, 11, 13).

Integration with the VANISH cohort demonstrated that NF induced pDARs correspond to transcriptional states linked to clinical outcomes (35). Because chromatin accessibility changes often precede and constrain transcription, these regulatory events likely occur early in the inflammatory cascade (35). The observed concordance between accessibility and RNA expression supports a model in which neutrophil chromatin remodeling reflects upstream regulatory mechanisms during sepsis. This finding is relevant with NET-associated disease, since NET release can amplify inflammation through the release of damage-associated molecular patterns (DAMPs) reinforcing a positive feedback loop of immune activation (6, 7, 49). Targeting upstream chromatin regulatory pathways could provide opportunities to disrupt this cycle before further immune dysregulation.

Clinical sepsis is increasingly recognized as heterogeneous comprised of multiple immune endotypes (11, 12, 16). More recently, studies including SUBSPACE (15) PANTHER (48) and ImmunoSep (47) emphasize the need to observe patients based on the immunobiology rather than uniform therapeutic strategies (47) .

Although integration with the SUBSPACE cohort (15) was exploratory and limited by gene overlap, it identified a small set of genes whose expression tracked with sepsis disease progression. Rather than directly identifying diagnostic biomarkers, this work illustrates how physiologically relevant stimulation can connect chromatin accessibility to clinical immune endotypes. The association of GM-CSF and Combo stimulation further indicates that inflammatory signaling environments may prime neutrophils toward specific transcriptional states associated with disease progression, suggesting the importance of upstream regulatory contexts over single gene effects.

The prominence of GM-CSF and STAT associated accessibility programs in severity linked genes is consistent with emerging precision medicine strategies that aim to match immune therapies, including JAK/STAT pathway inhibitors (such as baricitinib) (48) to patients based on underlying immune states and cytokine profiles. Collectively, this work establishes a mechanistic bridge between circulating inflammatory signaling, transcriptional endotypes, and clinical outcome, offering a way to refine patient stratification in sepsis and other NET-associated inflammatory diseases.

While we show a connection between our accessibility and existing expression data, a limitation was that the integration with clinical datasets was exploratory and constrained by cohort design and gene overlap. As a result, ROC analyses were not designed to establish diagnostic performance. Despite these limitations, the consistency observed across independent datasets supports the relevance of NF associated chromatin accessibility changes in neutrophils to clinically relevant inflammatory states. Collectively, these findings establish a foundation for future studies linking chromatin to precision therapeutic strategies in sepsis and other NET-associated inflammatory diseases.

Treating neutrophils with physiologically relevant factors in the context of whole blood enabled controlled stimulation of whole blood with defined inflammatory activators while preserving chromatin states for downstream profiling. This extends understanding of neutrophil responses to known immune activators in inflammatory disease. However, we cannot conclude whether the accessibility changes result solely from NFs acting on neutrophils or from secondary signaling cascades from other blood cell types, which then impact neutrophils.

Despite decades of study, therapeutic strategies targeting sepsis have largely failed to improve outcomes (50). One contributing factor could be the challenges of capturing individual immune variability and the complex inflammatory signaling in patients. Uniform stimulation and simplified *in vitro* systems likely obscure the regulatory heterogeneity that informs therapeutic responsiveness. Our approach of whole blood NF stimulation exposes donor dependent chromatin responses that could help inform future therapies. This framework supports shifting experimental models toward approaches that better incorporate immune diversity, enabling identification of regulatory pathways relevant for precision medicine-based approaches.

## Data availability statement

The original contributions presented in the study are publicly available. This data has been submitted to BioProject ID SUB15939056.

## Author Contributions

JC and TKK conceived the study, JC and TKK designed the experiments. BA performed the experiments, JC and AST provided guidance. JC and BPB performed data interpretation. All authors participated in the interpretation of data. JC, TKK, and AR wrote the manuscript, and all authors edited the manuscript for important intellectual content.

## Acknowledgments

We would like to thank Christina Wheeler and Finley Serneo, as well as the broader team at VolitionRx for helpful review of the data and discussion. ChatGPT version 5.2 was used to support the minor edits of R scripts and edits throughout the manuscript.

## Supplementary Tables

1. Donor Information
2. Stimulation Conditions
3. Illumina Sequencing Statistics
4. PMA and Combo Differential Regions
5. Natural Factor Differential Regions
6. pDAR and VANISH Concordance

