## Supplementary Figures for "Cytokine-induced Chromatin Accessibility in Whole Blood Neutrophils Links to Sepsis Transcriptional States"

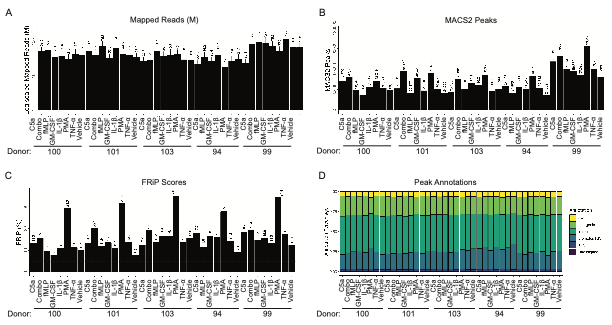

**Supplementary Figure 1**:

A. Total number of merged mapped reads for each of the donors, treatment, and time (x-axis). Total number of mapped reads (in millions) is on the y-axis. Dashed red line (y=10) indicates sequencing depth threshold.
B. Similar to (A) but with total number of MACS2 (q < 0.01) peaks.
C. Similar to (A) but FRiP percent is shown on the y-axis. Dashed red line (y=5) indicates FRiP threshold requirement.
D. Peak annotations generated on called MACS2 peaks (from (C)) for the samples shown as a percentage. (Transcription start site (TSS) Transcript termination sites (TTS)).

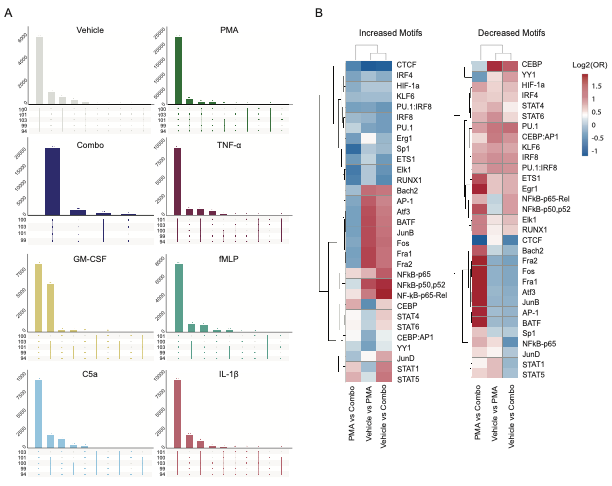

**Supplementary Figure 2**:

A. UpSetR plots showing the peak overlaps between donors across stimulation from Figure 1B-1C (grey: vehicle, dark blue: Combo, dark red: TNF-α, yellow: GM-CSF, light blue: C5a, green: fMLP, light red: IL-1B).
B. HOMER motifs within the significant DARs in each comparison from Figure 2E (Left: vehicle vs PMA, middle: vehicle vs Combo, right: PMA vs Combo). Results are plotted by -log10(p.value) (y-axis) and HOMER rank (x-axis). Increased motifs are indicated in red (right side of individual panel) and decreased motifs are indicated in blue (left side of individual panel).

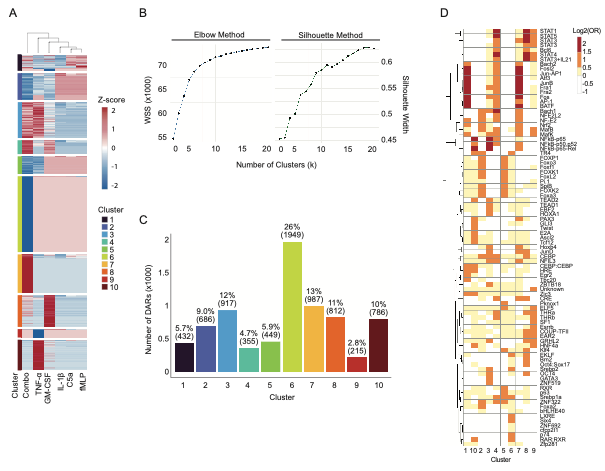

**Supplementary Figure 3**:

A. Unsupervised hierarchical clustering heatmap (k=10) on DAR composite activation score and then Cluster Signal was used to visualize (mean signal across entire cluster) across all stimulations (x-axis) and assigned cluster (y-axis). Columns were hierarchical clustered.
B. Clustering determination using Elbow Method (left) and Silhouette Method (Right).
C. Total number of DARs across all clusters (k=10). For cluster assignment colors: 1 – dark blue, 2 - blue, 3 - light blue, 4 – teal, 5 – green, 6 – light green, 7 – light orange, 8 – orange, 9 – red, 10 – dark red.
D. Full Heatmap of HOMER Motif analysis across each cluster (y-axis). Cluster number is indicated on the x-axis and Rows are the TF motif. Color indicates increasing fold enrichment from log2(Odds Ratio) (white: -1-0, yellow: 0-0.5, orange: 0.5-1, red: 1.5-3).

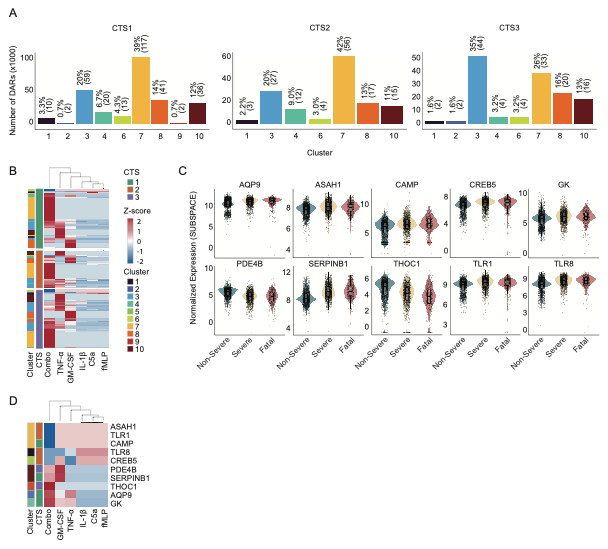

**Supplementary Figure 4**:

A. Distribution of ATAC cluster assignments amongst concordant promoter-DARs for each CTS module separately. CTS1 (left), CTS2 (middle), and CTS3 (right) are shown. The x-axis shows the number of regions assigned to each chromatin cluster, highlighting shared and distinct regulatory composition. For cluster assignment colors: 1 – dark blue, 2 - blue, 3 - light blue, 4 – teal, 5 – green, 6 – light green, 7 – light orange, 8 – orange, 9 – red, 10 – dark red.
B. Z-score normalized peak counts across the top concordant promoter DARs from the three CTS modules (green: CTS1, orange: CTS2, purple: CTS3) across the clusters (1 – dark blue, 2 - blue, 3 - light blue, 4 – teal, 5 – green, 6 – light green, 7 – light orange, 8 – orange, 9 – red, 10 – dark red.). Stimulation conditions (column) and DARs (rows) were hierarchical clustered.
C. Expression of ATAC-linked genes in the SUBSPACE cohort across sepsis severity grades (Green – non-severe, yellow – severe, and red – fatal). Violin and boxplots show the distribution of normalized gene expression for the ten overlapping genes (*AQP9, ASAH1, CAMP, CREB5, GK, PDE4B*, *SERPINB1*, *THOC1*, *TLR1*, and *TLR8*). Individual points represent individual donors (non-severe: n=1,293, severe: n=1,203, and fatal: n=387).
D. Heatmap of promoter-associated chromatin accessibility for the ten genes across all stimulation conditions. Rows represent the promoter-associated peak per gene, and columns represent the stimulation compared to vehicle. Values are z-score normalized across the comparisons. Row annotations indicate ATAC cluster assignment and CTS module (CTS1 – green, CTS2 – orange, CTS3 – purple).
